# Open-land-derived agroforestry and effects of abandonment of management of the main crop on ecosystem services and woody plant diversity

**DOI:** 10.1101/2025.05.27.656282

**Authors:** Lucas M. Fonzaghi, Kristoffer Hylander, Thiago S. de Melo, Ayco J. M. Tack

**Affiliations:** Department of Ecology, Environment and Plant Sciences, Stockholm University, Stockholm, SE-106 91, Sweden; Universidade do Oeste Paulista, Presidente Prudente, BR-19050-920 São Paulo, Brasil

**Keywords:** abandonment, agroforestry, Atlantic forest, *Coffea arabica*, conservation, ecosystem services, ecosystem disservices

## Abstract

Tropical forests are biodiversity hotspots threatened by deforestation and fragmentation. One strategy to restore tropical forest is to establish agroforestry on open land. However, we lack insights into the social and ecological dynamics of open-land-derived agroforests, and specifically the reasons for and consequences of abandonment. We combined semi-quantitative interviews with biodiversity surveys to examine the reasons for abandonment and the consequences for ecosystem services, disservices, and the woody plant community in the Atlantic Forest region. Reasons for abandonment were primarily related to agroecology. Ecosystem services, disservices, and woody plant diversity had not (yet) differentiated after the abandonment of management. Strengthening agronomic support to farmers is essential to maintain and expand open-land-derived agroforestry. Notably, as abandoned agroforests were not converted back to open land, partly due to legal restrictions, they continued to provide ecosystem services and contributed to the restoration of native tree cover across the landscape.

## Introduction

Tropical forests are among the most biodiverse ecosystems in the world but are rapidly declining due to deforestation (Brancalion et al., 2019). To reverse the loss of tropical forests, biodiversity and ecosystem services, several conservation initiatives have been launched (Brancalion et al., 2019; Joly et al., 2014). These initiatives include active restoration areas where native trees are planted under strict conservation laws, ecological corridors, and use of native trees in agroforestry (Giudice Badari et al., 2020; Valladares-Padua et al., 2002). While agroforestry might degrade the environment when established in old-growth forests through thinning (Martin et al., 2020), when established on degraded open land, agroforestry has a major potential to recover native biodiversity and ecosystem services (Giudice Badari et al., 2020; Martin et al., 2020; Shennan-Farpón et al., 2022). Indeed, the restoration potential of open-land-derived agroforests has been shown to match active restoration areas in terms of richness and tree density (Giudice Badari et al., 2020), and in some cases, the agroforests can even match old-growth forests in aboveground biomass and canopy cover (Giudice Badari et al., 2020). Open-land-derived agroforests can also increase natural resources for different providers of ecosystem services, such as pollinating insects, and create stepping stones for biodiversity between larger nature reserves (Martin et al., 2020; Schuler et al., 2022; Shennan-Farpón et al., 2022). However, while forest-derived agroforestry systems are quite well understood, we lack insights into the social and ecological dynamics of open-land-derived agroforests (Martin et al., 2020).

Beyond ecosystem restoration, agroforests need to be beneficial to farmers. While it is not uncommon for farmers to receive support for or during the establishment of an agroforest, the long-term management depends on the agroforest continuing to be beneficial to the farmer. This in turn depends on the balance between benefits and disbenefits related to agroforestry. The majority of agroforests are based on one or a few main crops, such as coffee, cacao and vanilla, but can simultaneously provide other provisioning services such as fruits, medicinal herbs, and wood for fuel or tools (Martin et al., 2020; Schuler et al., 2022; Shennan-Farpón et al., 2022). In addition to the provisioning ecosystem services, agroforests provide regulating services such as pollination, seed dispersal, improvement of soil quality, shade, buffering of high and low temperatures, protection against drought, and shelter against rain and wind (Martin et al., 2020; Schuler et al., 2022; Shennan-Farpón et al., 2022). Despite the diverse set of provisioning and regulating ecosystem services, agroforests can also introduce disservices to the farmers, such as providing shelter for crop raiding animals, dangerous predators, venomous snakes and arachnids. An alternative reason for adoption of agroforests is to follow state regulations. In some countries, conservation laws stipulate that farmers must have a certain amount of land area covered by native trees, and in this case farmers can convert open land to agroforestry to comply with the law (Martin et al., 2020; Schuler et al., 2022; Shennan-Farpón et al., 2022). From the perspective of establishing a successful agroforestry system, several stages are important. Regarding the initial adoption of agroforestry, there are several barriers, including financial costs, limited knowledge on agroforestry, labour requirements, and lack of governmental and non-governmental support (Shennan-Farpón et al., 2022). Yet, even after the initial adoption of agroforestry and planting of the main crop(s) and shade trees, several challenges could lead farmers to abandon management of the main crop (Fig. 1A; Rolim et al., 2017). However, the processes leading to the abandonment or continued management of agroforests are rarely studied (but see Aragón et al., 2021; Arnold et al., 2021; Lozada et al., 2007; Richter et al., 2007; Rolim et al., 2017). We propose that in the short term, the main crop and shade trees could fail to establish, after which the farmers could either retry to establish the agroforest or repurpose the land (Fig. 1A). In the longer term, the crop might drop in productivity, the crop might lose productivity altogether (Giudice Badari et al., 2020), or unrelated socioeconomic problems occur (Richter et al., 2007; Rolim et al., 2017; Shennan-Farpón et al., 2022), which could all lead to abandonment of management of the main crop (Fig. 1A). However, we lack insights into the factors that result in abandonment of management of the main crop, especially in open-land-derived agroforestry.

**Figure 1:**
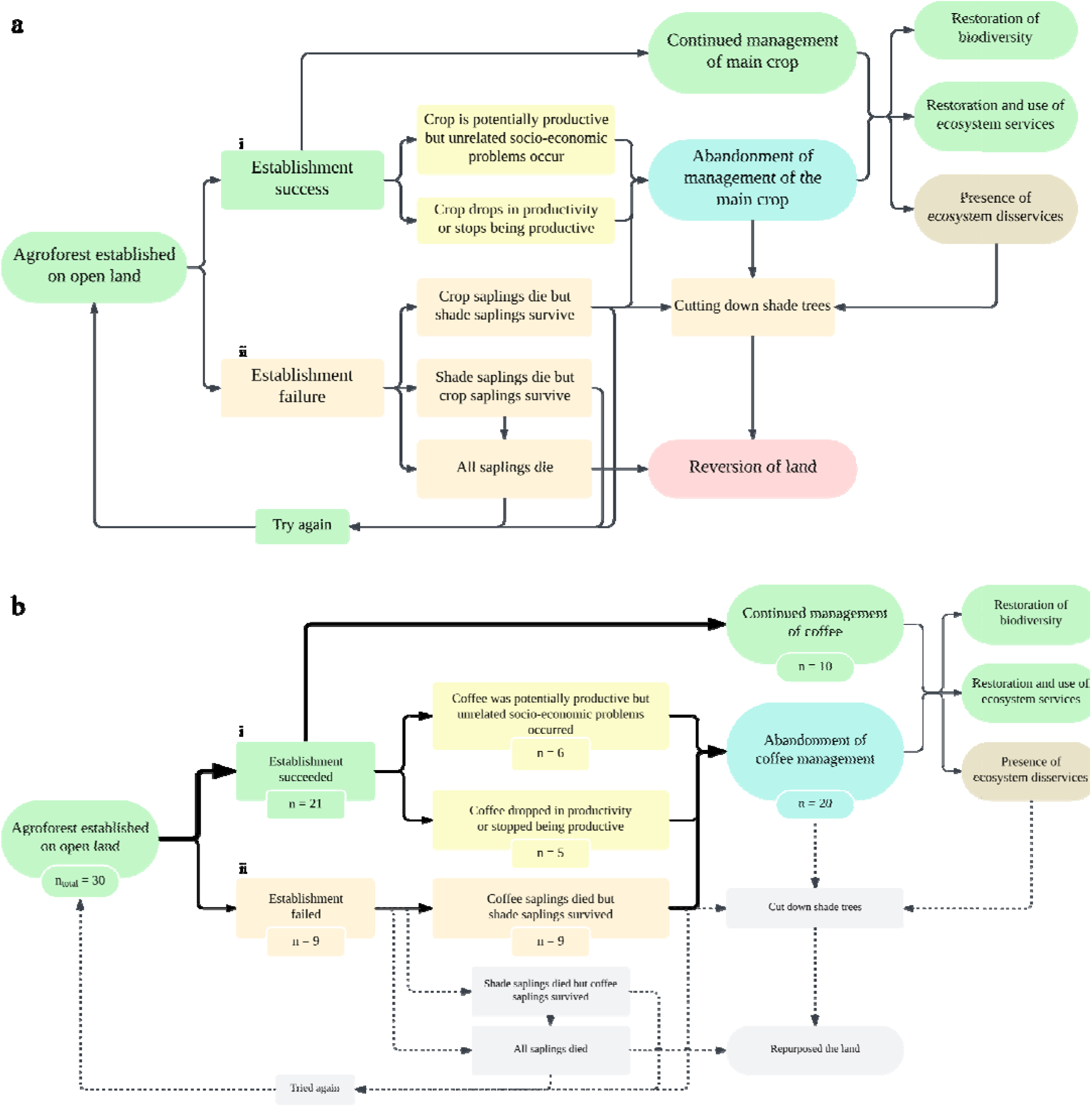
Schematic overview of the (a) potential and (b) observed processes and outcomes of open land conversion to agroforests. Panel (a) illustrates two potential outcomes of the initial establishment: (i) agroforest establishment success or (ii) establishment failure due to crop or shade tree death. If the crop fails, the farmer may retry, or shade trees might establish without the crop. A successful initial establishment can lead to continued management, though socioeconomic problems or productivity drops may still cause abandonment of the management of the main crop. Even in the case of abandonment, native shade trees can still provide ecosystem services and disservices and contribute to conservation. Abandonment and increased ecosystem disservices might also result in removal of the native trees and reversion to open land. Panel (b) shows how the 30 investigated coffee agroforestry sites in *Pontal do Paranapenama* developed over time, using the conceptualization from panel (a). Arrow thickness reflects the prevalence of the pathway, and dotted lines and greyed boxes represent processes and outcomes that were not observed in our study.

Beyond the reasons for abandonment of the main crop, we lack insight into how open-land-derived agroforestry develops in terms of structure and composition of the tree community, and how this depends on the abandonment of management of the main crop. For example, it could be that agroforests where the management of the main crop is abandoned contribute to the same extent to the conservation of native forests as agroforests where the management of the main crop is maintained. However, there could also be differences between the two types of sites. A few studies have compared abandoned and managed sites within forest-derived agroforest systems, where abandonment generally increased species richness and structural complexity (Aragón et al., 2021; Arnold et al., 2021; Lozada et al., 2007; Rolim et al., 2017). For example, after 40 years of abandonment of cocoa agroforests in Bahia (northeast Brazil), abandoned cocoa agroforests had higher species richness than managed agroforests (Rolim et al., 2017), and species richness and structural complexity increased with time after abandonment of cocoa agroforests established 80 years ago in Trinidad and Tobago (Arnold et al., 2021). Other examples include increased diversity of saplings (Lozada et al., 2007) and increased abundance of insects (Richter et al., 2007) in abandoned forest-derived coffee agroforests when compared to still managed coffee agroforests. However, no studies have investigated the ecological effects of abandonment in open-land-derived coffee agroforests. Understanding how open-land-derived agroforests develop, both when maintained and when the management of the main crop is abandoned, is crucial for the development of conservation policies and actions to reverse the decline of tropical forest.

A key biodiversity hotspot within the tropical forest region is the Brazilian Atlantic Forest. The Brazilian Atlantic Forest is severely threatened by historical and on-going deforestation, and currently only 10% of the original cover remains (Joly et al., 2014; but see Rezende et al., 2018). The Atlantic Forest has been the main target of multiple restoration pledges by the Brazilian government, and several initiatives have aimed to establish agroforestry on open land to create ecological stepping stones (Joly et al., 2014; Lemos et al., 2021).

Our overarching aim was to improve knowledge on the reasons that lead to the abandonment of management of open-land-derived agroforests, and the effects of this abandonment on the presence of ecosystems services and disservices, as well as on the structure of the woody plant community. For this, we focused on the Brazilian Atlantic Forest of the northwest São Paulo region of Pontal do Paranapanema and selected 30 smallholders who had established coffee agroforests on open land as part of local programs. We conducted semi-quantitative interviews with the farmers and surveyed their agroforests to answer the following questions:

1. What are the reasons that sometimes lead to the abandonment of management of the main crop within agroforests?
2. Does the abandonment of management of the main crop change the presence of ecosystem services and disservices when compared to agroforests that are still managed?
3. What are the differences in the diversity and structure of the woody plant community between agroforests that are still managed and agroforests where management of the main crop was abandoned?

## Material and Methods

### Study system

The region Pontal do Paranapanema is located in the northwest of São Paulo state, Brazil, between the Paraná and Paranapanema rivers (Fig. S1). The region is around 19,000 km^2^ and has an elevation ranging from 265 to 320 m (Giudice Badari et al., 2020; Rossi, 2017; Valladares-Padua et al., 2002). The region has a mean annual temperature of 24°C, and a mean annual rainfall of 1341 mm, and the soils are predominantly red latosols and red argisols, which are sandy soils prone to erosion (Giudice Badari et al., 2020; Rossi, 2017; Valladares-Padua et al., 2002). The native vegetation belongs to the semi-deciduous type of the Atlantic Forest domain (Valladares-Padua et al., 2002). Today, only fragments of the local vegetation remain, with the largest remnant being the Morro do Diabo State Park (shaded area in Fig. S1), which is a major habitat for one of the last populations of the jaguar *Panthera onca* in this region (Giudice Badari et al., 2020; Valladares-Padua et al., 2002). Pontal do Paranapanema has been a site of land disputes in the 1990s involving the Landless Worker Movement (MST) who occupied unproductive land (Valladares-Padua et al., 2002). Today, the region is mostly used for pastures, sugarcane production, and small-scale farming (Giudice Badari et al., 2020; Valladares-Padua et al., 2002). Laws at federal level stipulated that agricultural properties are required to have 20% of their area covered with native vegetation (Law No. 12,651, 2012). As natural forest covers only c. 10% of the landscape, establishing open-land-derived forests and agroforests was a promising option to increase the coverage with native vegetation and thereby comply with the law (Martin et al., 2020). Between 2005 and 2020, a local NGO (Instituto de Pesquisas Ecológicas, often shortened to IPÊ) has helped local smallholder farmers to establish Arabica coffee-based agroforests on open land by planting coffee and shade trees native to the semi-deciduous vegetation of the Atlantic Forest. Some of the farmers also included fruit trees such as citrus and avocado. The native shade trees were planted in rows with a gap of 6 m to 8 m, intercalated with rows of coffee and occasionally fruit trees. Subscription to the agroforestry program was voluntary, and there were limited spots available (c. 50 across the years).

### Site selection

To select the sites, we used an initial list with 36 names of farmers that joined IPÊ’s agroforest initiative. Their farms were unmapped but located within known MST settlements in the municipalities of *Mirante do Paranapanema, Teodoro Sampaio*, and *Euclides da Cunha Paulista*. We located the farms by asking locals within each settlement where to find the farmers on the list. The final 30 sites were selected based on which farms were found and which farmers agreed to participate in the study (Fig. S1). During site selection, we only encountered a single farmer from the list on whose site no shade trees were present; the reason provided was fire, and the farmer did not agree to take part in a longer interview. The farmers had settled their lands between 9 and 44 years ago and established the agroforests between 7 to 23 years ago (Fig. S2). The size of the farms ranged from 12 to 32 ha, and agroforests covered 1 to 2 ha. Ten agroforests were still productive and managed for coffee, whereas coffee management had been abandoned in twenty agroforests between 3 and 16 years ago (Fig. S3).

### Interviews

To understand the dynamics of establishment, the mechanisms of abandonment and the perceptions on ecosystem services and disservices we conducted semi-quantitative interviews with the farmers. The questions focused on (i) when the agroforest was established (henceforth ‘time since establishment’), (ii) if relevant, the reasons for abandonment of coffee management and when abandonment took place (henceforth ‘time since abandonment’); and (iii) the perception of ecosystem services and disservices. For the full set of questions, see appendix A.

### Biodiversity survey

To examine the structure of the woody plant community in the agroforests, we recorded the occurrences of woody plant species in a sample area of 10 × 10 m in the centre of the first established agroforest area of the farm. We identified the species and recorded the diameter at breast height (DBH) of all woody plant specimens which had DBH > 1 cm. Specimens which were not identified *in situ* were sampled and photographed for later identification. Samples were pressed and stored in the herbarium of the Federal University of São Paulo. For the full list of recorded species and their abundance, see Table S1. We also counted any woody plants which were larger than 1.5 m but had a DBH < 1 cm, without identifying them. We took five pictures of the canopy from above the coffee shrub layer at each corner and at the centre of the sample area and used the ImageJ plugin Hemispherical 2.0 to calculate canopy cover (Beckschäfer, 2015).

### Data analysis

All of the analyses were done in R v. 4.4.

#### Reasons FOR ABANDONMENT OF MANAGEMENT OF THE MAIN CROP

To understand the reasons that led to the abandonment of coffee management, we described and visualized the distribution of answers related to this interview question.

#### Ecosystem SERVICES AND DISSERVICES IN MANAGED AND ABANDONED AGROFORESTS

To examine if abandonment of management affected the presence of ecosystem services and disservices, we tested if there were significant differences in the proportions of ‘yes’ and ‘no’ answers to the interview questions related to the perception of ecosystem services and disservices between farmers who still manage their main crop and farmers who abandoned management. For this, we used a Fisher’s exact test (Quinn & Keough, 2023).

#### The EFFECTS OF ABANDONMENT ON THE STRUCTURE OF THE WOODY PLANT COMMUNITY

To understand how abandonment of the management of the main crop affects the structure of the woody plant community in agroforests, we focused on six descriptors of diversity and structure: (i) species richness, (ii) Shannon’s diversity index, as calculated using the function *diversity* from the *vegan* package (Oksanen et al., 2025), (iii) abundance of woody plants with DBH larger than 1 cm, (iv) abundance of woody plants with DBH smaller than 1 cm, (v) Simpson’s evenness (calculated by dividing one over Simpson’s diversity index by the species richness) and (vi) canopy cover. We then modelled each metric as a function of abandonment (‘managed’ or ‘abandoned’), time since establishment, time since abandonment and the interaction between time variables using the functions *lm* and *glm* in base R. Shannon’s diversity index was normally distributed, evenness was normally distributed after log-transformation. We used the *Poisson* family for richness, and *quasipoisson* family for the abundance variables and for canopy cover. We used backward model selection with a threshold of P < 0.1 to retain predictor variables to reach a final model (Crawley, 2007) and assessed significance using the function *Anova* in the package *car* (Fox et al., 2024). Model assumptions, such as normality of residuals and equal variance, were visually evaluated, and no significant correlations among the predictor variables were found by using the *cor* function (Crawley, 2007).

## Results

### Reasons for abandonment of management of the main crop

Management of coffee was abandoned in twenty of the thirty agroforests (Fig. 1b). Abandonment was mainly related to agroecological reasons, including failure of establishment of coffee saplings (n = 9) or a decline in productivity (n = 5) (Fig. 1b and S4). Six farmers abandoned the management of coffee due to socio-economic reasons unrelated to the productivity of the coffee, such as lack of time, health issues or financial failure (Fig. 1b and S4).

### Ecosystem services and disservices in managed and abandoned agroforests

Only one out of the 14 ecosystem services differed significantly between managed and abandoned agroforests, whereas none of the ecosystem disservices differed between managed and abandoned agroforests (Fig. 2). The only ecosystem service that significantly differed between managed and abandoned agroforests was timber, which was collected more often by farmers who still managed their agroforest (Fig. 2 and Table S2). Moreover, there was a trend for farmers who still managed their agroforests to report more often that they feel their agroforest is aesthetically pleasing and report more often the large vertebrates such as tapirs, black tamarins and jaguars (Fig. 2 and Table S2).

**Figure 2:**
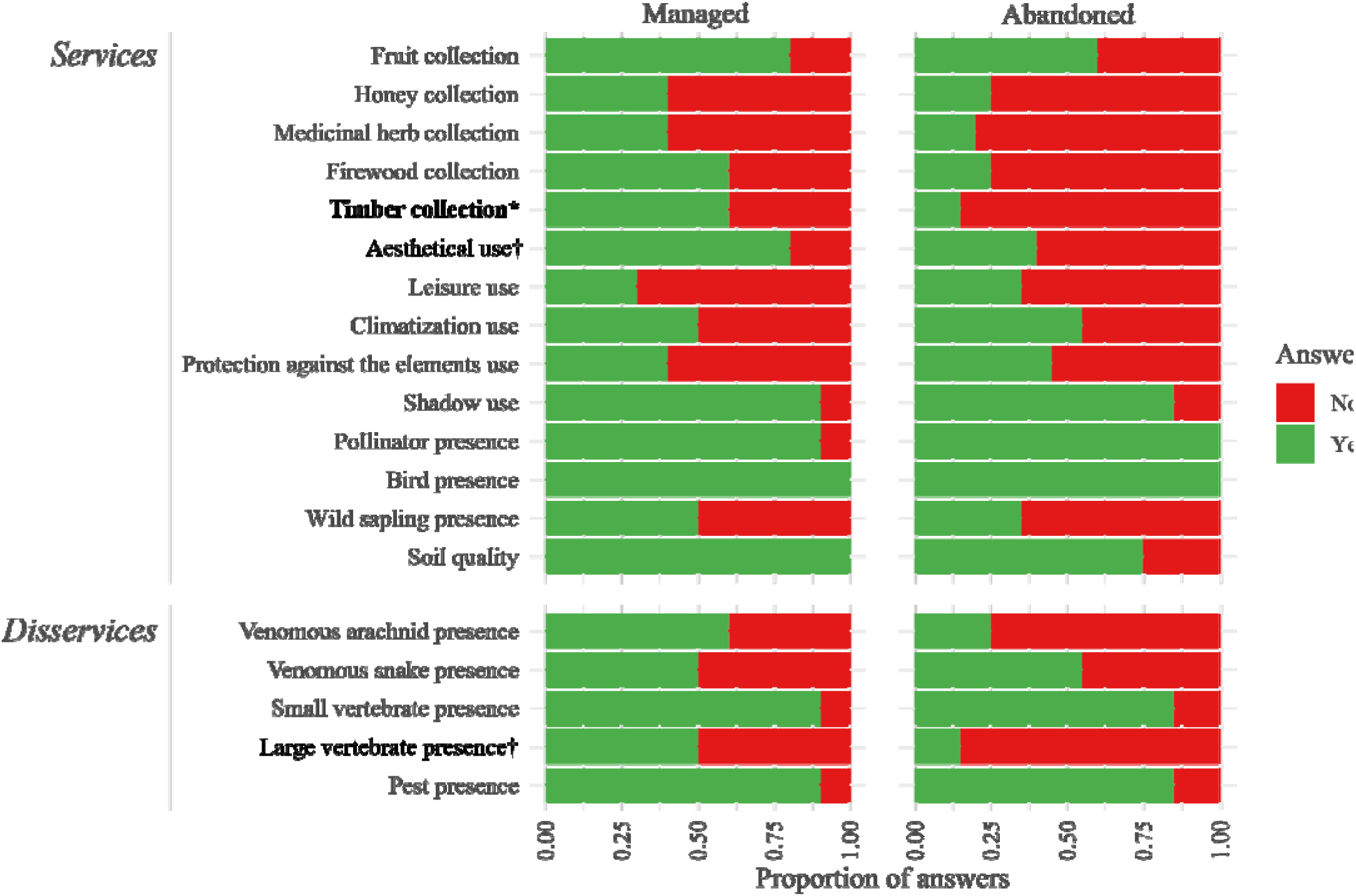
Ecosystem services and disservices in managed and abandoned agroforests. Shown is the proportion of yes (green) and no (red) answers given by farmers to questions related to the presence of ecosystem services and disservices, separately for farmers who managed and abandoned coffee management. Ecosystem services and disservices that differed significantly (P < 0.05) between managed and abandoned agroforests are in bold with an asterisk, and those for which there was a trend (P < 0.1) are marked with a dagger sign (†). For detailed statistics, see Table S2.

### The effects of abandonment on the structure of the woody plant community

The species richness, Shannon diversity, abundance, evenness and canopy cover did not differ between managed and abandoned agroforests (Fig. 3 and Table S3). Likewise, time since establishment and time since abandonment did not affect most of the descriptors of the woody plant community (Table S3) with the exception of a weak effect of time since establishment on canopy cover in agroforests that were abandoned for more than 5 years, though this could be explained by outliers in the data (Fig. S5).

**Figure 3:**
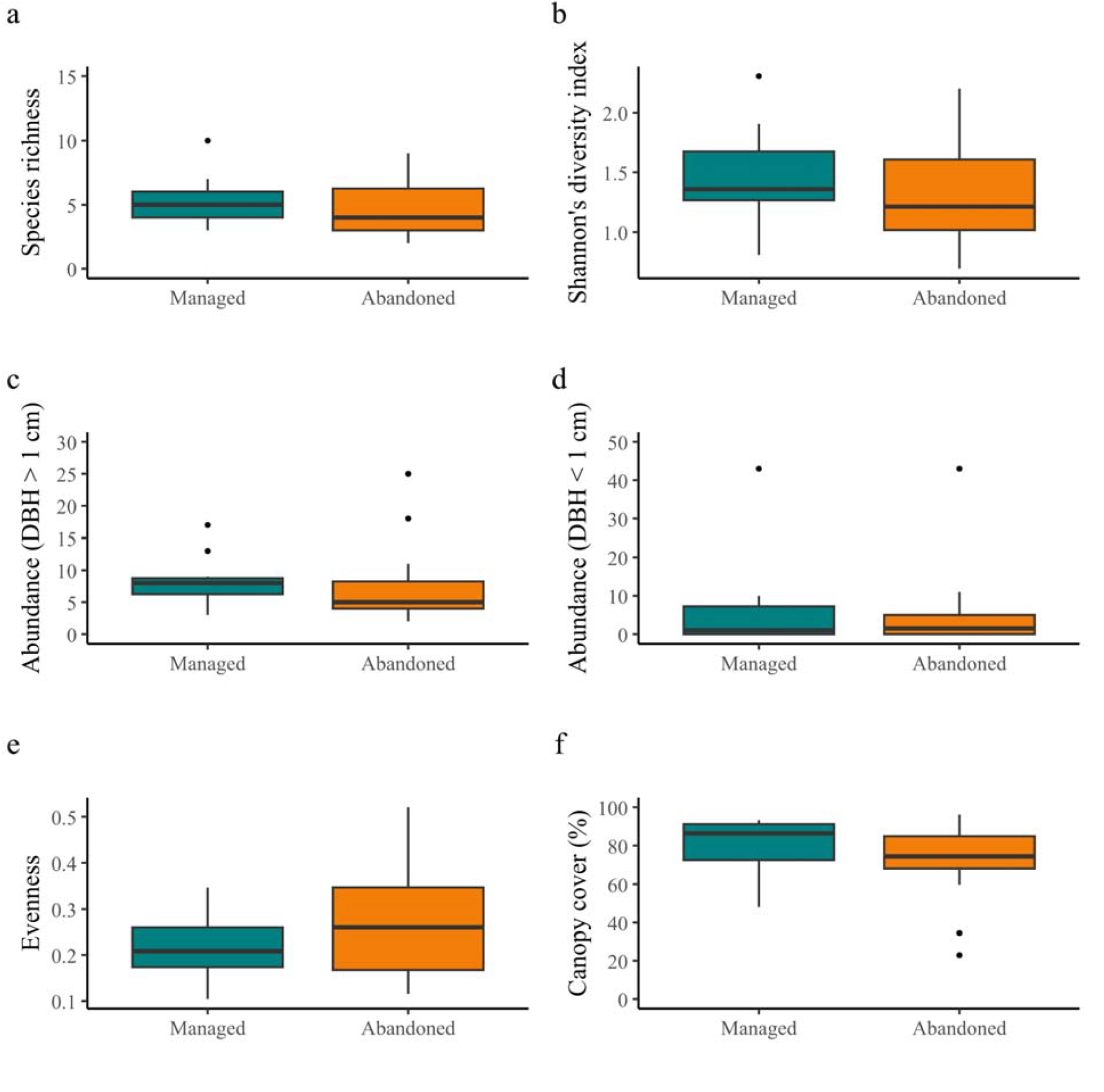
The structure of the woody plant community in managed (N = 10) and abandoned (N = 20) agroforests. Shown are box plots. Presented are (a) species richness, (b) Shannon’s diversity index, (c) abundance of woody plants with DBH larger than 1 cm, (d) abundance of woody plants with DBH smaller than 1 cm, (e) evenness and (f) canopy cover. Turquoise represents agroforests which are still managed, and orange represent agroforests in which coffee management was abandoned. Boxes represent the interquartile range, the black line represents the median, whiskers extend to data points up to 1.5 times the interquartile range and individual dots are outliers. For detailed statistics, see Table S3.

## Discussion

The overarching aim of our study was to understand the mechanisms and consequences of the abandonment of management of the main crop, coffee in our case, in open-land-derived agroforests. We found that abandonment was common (two-third of sites), and the major reasons for abandonment were related to agroecology, whereas only few farmers gave socioeconomic reasons. Farmers from managed and abandoned sites perceived mostly the same services and disservices from their agroforests. The abandonment of coffee management had no significant effect on the structure of the woody plant community. The low success rate of establishment of agroforestry on open land due to agroecological challenges might be alleviated by more and continued external support. Importantly, even if many agroforests were unsuccessful in terms of coffee production, the farmers perceived other services from having the agroforest on their land, and the farmer movement complied with nature conservation laws by increasing forest cover in the landscape. At the same time, the agroforests support the conservation of native trees and their associated biodiversity, one of the aims of IPÊ’s project in the region.

### Reasons for abandonment of management of the main crop

In our study area in Pontal do Paranapanema, we found that two-thirds of the farmers had abandoned coffee management. Studies conducted in abandoned agroforests in Brazil and other tropical regions have rarely investigated the reasons for abandonment. Some reasons that have been given in published studies where agroforests were found to be abandoned are changes in market prices (Aragón et al., 2021; Arnold et al., 2021; Lozada et al., 2007; Richter et al., 2007; Rolim et al., 2017), mismanagement (Aragón et al., 2021), rural emigration (Aragón et al., 2021), and competition (Arnold et al., 2021). However, none of these studies asked the farmers in their study area about their reasons for abandonment, but rather cited a secondary reference or did not provide any reference. In our study, we found that the majority of farmers abandoned coffee management due to failure of establishment of the shrubs and declines in productivity, whereas some farmers cited reasons unrelated to the productivity of the coffee, such as lack of time, health issues, change in ownership or financial problems. Hence, initiatives to prevent abandonment of management should not only support farmers more during the initial phase of establishment of the coffee saplings but also provide continued support to maintain crop productivity. As coffee agroforestry is relatively new to the study area, it is crucial to develop agricultural techniques and management guidelines tailored to local farmers.

While not the focus of our study, most farmers mentioned drought as a reason for the failure of initial establishment of coffee saplings, and farmers who successfully established their agroforests reported the lack of permission to prune the native trees and the resulting high shade levels as a reason for the decrease in productivity of the coffee. This reasoning matches previous studies which have shown that the photosynthetic rate of coffee plants decreases when shade levels are too high (Getachew et al., 2022). Despite the failure of establishment, the abandonment of coffee management did not imply removal of shade trees, as farmers face legal restrictions and possible fines for cutting native trees. Importantly however, the great majority of farmers did not regret adopting agroforestry (see also next section). At the same time, none of the farmers tried to re-establish coffee production within the created agroforests, or expressed any wish to expand their agroforestry areas, which is not surprising given the lack of establishment success, and the feeling that there was a need for more support from local governmental and non-governmental organizations. These findings corroborate the results of a study by Shennan-Farpón et al. (2022), conducted in the same region, that found that the most important barriers to the adoption of agroforestry were the lack of support from the government and local community organisations (Shennan-Farpón et al., 2022).

### Ecosystem services and disservices in managed and abandoned agroforests

The perception of the presence of ecosystem services and disservices was very similar in managed and abandoned agroforests. The most likely reason seems to be the congruent similarity in the diversity and structure of the woody plant community in managed and abandoned agroforests (see next section). Hence, the shade trees benefited the farmers with provisioning and regulating ecosystem services, irrespective of whether they managed the agroforest for coffee. There is a lack of comparable data on how ecosystem services are perceived by farmers who abandoned management as compared to those who still maintain management, be it open-land or forest-derived agroforestry. However, one study comparing farmers with and without agroforests found that farmers who established open-land-derived agroforests reported the presence of ecosystem services (disservices are not reported) more often than farmers in the same region who chose not to establish agroforests (Shennan-Farpón et al., 2022). Taken together with our own findings, this could mean that having the (agro)forest on the farm is enough to result in an increase in ecosystem services, and management of the crop in the agroforest is not necessary for ecosystem services to be present. The few differences in perception of ecosystem services and disservices between farmers who abandoned management and those who still maintain management in our study could be due to the frequency of farmer visits to the agroforests: Farmers who still manage their agroforest will generally visit the agroforest more frequently, and thereby have more opportunities to collect firewood, enjoy the aesthetic value of the native trees and observe organisms such as large vertebrates. Interestingly, the most reported disservice was the presence of *saúva* leafcutter ants (*Atta* sp., reported as pests in our results), which attack both the farmers’ vegetable gardens and the native trees in the agroforests. We suspect that this may be one of the reasons for the failure of establishment of coffee at earlier stages, and we recommend that stakeholders take the potential presence of leafcutter ants into account when establishing future open-land-derived agroforestry in the region. The most reported disservices in our study (leafcutter ants and small vertebrates such as invasive European hares) were of a different type than the ones reported in coffee agroforests in Ethiopia, in which large vertebrates such as wild pigs and monkeys were considered the main disservice to crops (Ango et al., 2014). Importantly, farmers perceived the balance between ecosystem services and disservices to be in favour of the services.

### The effects of abandonment on the structure of the woody plant community

The diversity and structure of the woody plant community was very similar between managed and abandoned agroforests. As such, our study shows that abandonment does neither have a positive nor negative impact on the diversity and structure of the woody plant community on open-land-derived agroforests. While we did not find other studies that focused on comparing the woody plant community in managed and abandoned open-land-derived agroforests, we found three studies comparing the woody plant community in managed and abandoned forest-derived agroforests, which reported a higher diversity in abandoned agroforests. We propose three hypotheses for the differences between our study and previous studies. First, previous studies were conducted in forest-derived agroforests, and processes of forest regeneration after abandonment might differ between forest-derived and open-land-derived agroforests. Second, the timescale of our study was – with agroforests being abandoned 3 to 16 years ago – relatively short, and we might therefore expect differences to appear during the coming decades. The three studies comparing the woody plant community in managed and abandoned forest-derived agroforests support the hypothesis that time since abandonment plays a major role in understanding the differences between managed and abandoned agroforests. Two studies focusing on relatively old abandoned agroforests (25 to 80 year old forest-derived abandoned cocoa agroforests in Trinidad and Tobago and 40 year old forest-derived cocoa agroforests in the northeastern Brazilian Atlantic Forest; Arnold et al., 2021; Rolim et al., 2017) reported that tree species richness, Shannon’s diversity index, and tree abundance were higher in abandoned agroforests and increased with time since abandonment. In contrast, in a study on coffee agroforests that were abandoned 10-15 years ago, adult tree species richness was similar between abandoned and managed agroforests, and only the species richness of saplings was higher in abandoned agroforests compared to managed agroforests (Lozada et al., 2007). While the time hypothesis is credible, we note that we did not detect a significant effect of time after abandonment within our current data set. The third reason for the similarity between the abandoned and managed agroforests in our study might be current management. While this was not a focus of the current study, we observed several times that farmers who abandoned management of coffee used the agroforest as shade for their cattle. These cattle in turn might trample seedlings, munch on young leaves, and break down young tree saplings, taking away the expected positive effect of abandonment on seedling regeneration. To distinguish between these three hypotheses, we hope that future studies will follow the development of the woody plant community in our study sites during coming decades, and either observationally or experimentally study how current management in the form of cattle grazing drives the development of the woody plant community through time in both abandoned and managed sites.

### Conclusion

We found that the abandonment of coffee management in the studied open-land-derived agroforests was mostly caused by agroecological problems, that farmers perceived ecosystem services and disservices irrespective of whether or not they managed the agroforest for coffee production, and that the composition and structure of the native tree community did not differ between managed and abandoned agroforests. The agroecological challenges faced by the farmers indicate that – if there is a wish for increased adoption or expansion of agroforestry – there is an urgent need for additional support of farmers by governmental and non-governmental organizations, during both establishment and later phases. Importantly, even though many agroforests were unsuccessful in terms of coffee production, the farmers perceived benefits from having the agroforest on their land and did not regret adopting agroforestry. As agroforests were not cut down after abandonment, they fulfilled one of the original aims of establishment: the restoration of native tree cover and biodiversity.

## Acknowledgements

We thank Aline Souza, our contact person at IPÊ (*Instituto de Pesquisas Ecológicas*) for providing the list of names of the farmers, Valter Ribeiro Campos for the help with species identification, and Carla Poleselli Bruniera and Isabella Capucho from the Federal University of São Paulo for help with identification and storage of samples in the university’s herbarium.

## Author contributions

Lucas Fonzaghi: Conceptualization, Methodology, Formal analysis, Investigation, Writing - Original Draft, Visualization, Project administration. Kristoffer Hylander: Conceptualization, Methodology, Writing – Review & Editing, Supervision. Thiago Sobral: Methodology, Investigation, Resources. Ayco Tack: Conceptualization, Methodology, Validation, Writing – Review & Editing, Supervision.

