## Supplemental Material for "Open-land-derived agroforestry and effects of abandonment of management of the main crop on ecosystem services and woody plant diversity"

**Table S1.** Tree and shrub species found in thirty agroforests in southeastern Brazil. Shown are the scientific species names, family, biogeographic status, and number of individuals found.

| **Species** | **Family** | **Biogeographic status** | **Number of individuals** |
| --- | --- | --- | --- |
| *Aegiphila integrifolia* (Jacq.) Moldenke | Lamiaceae | Native | 5 |
| *Albizia niopoides* (Spruce ex Benth.) Burkart | Fabaceae | Native | 1 |
| *Alchornea glandulosa* Poepp. & Endl. | Euphorbiaceae | Native | 2 |
| *Allophylus edulis* (A.St.-Hil. et al.) Hieron. Ex Nierderl. | Sapindaceae | Native | 6 |
| *Aloysia virgata* (Ruiz & Pav.) Juss. | Verbenaceae | Native | 3 |
| *Anadenanthera colubrina* (Vell.) Brenan | Fabaceae | Native | 3 |
| *Anadenanthera colubrina* var. *cebil* (Griseb.) Altschul | Fabaceae | Native | 1 |
| *Annona muricata* L. | Annonaceae | Cultivated | 3 |
| *Apuleia leiocarpa* (Vogel) J.F.Macbr. | Fabaceae | Native | 2 |
| *Astronium graveolens* Jacq. | Anacardiaceae | Native | 4 |
| *Astronium urundeuva* (M.Allemão) Engl. | Anacardiaceae | Native | 9 |
| *Cedrela fissilis* Vell. | Meliaceae | Native | 1 |
| *Ceiba speciosa* (A.St.-Hil.) Ravenna | Malvaceae | Native | 2 |
| *Cereus jamacaru* DC. | Cactaceae | Endemic | 1 |
| *Colubrina glandulosa* Perkins | Rhamnaceae | Native | 1 |
| *Copaifera langsdorffii* Desf. | Fabaceae | Native | 1 |
| *Cordia trichotoma* (Vell.) Arráb. Ex Steud. | Cordiaceae | Native | 1 |
| *Croton floribundus* Spreng. | Euphorbiaceae | Native | 7 |
| *Croton urucurana* Baill. | Euphorbiaceae | Native | 2 |
| *Dahlstedtia muehlbergiana* (Hassl.) M.J.Silva & A.M.G.Azevedo | Fabaceae | Native | 1 |
| *Delonix regia* (Bojer ex Hook.) Raf. | Fabaceae | Cultivated | 3 |
| *Dictyoloma vandellianum* A. Juss. | Rutaceae | Native | 11 |
| *Eugenia uniflora* L. | Myrtaceae | Native | 8 |
| *Gallesia integrifolia* (Spreng.) Harms | Phytolaccaceae | Endemic | 2 |
| *Genipa americana* L. | Rubiaceae | Native | 5 |
| *Gliricidia sepium* (Jacq.) Kunth ex Walp. | Fabaceae | Naturalized | 3 |
| *Guazuma ulmifolia* Lam. | Malvaceae | Native | 4 |
| *Handroanthus impetiginosus* (Mart. Ex DC.) Mattos | Bignoniaceae | Native | 11 |
| *Hymenaea courbaril* L. | Fabaceae | Native | 3 |
| *Inga edulis* Mart. | Fabaceae | Native | 5 |
| *Inga laurina* (Sw.) Willd. | Fabaceae | Native | 3 |
| *Inga marginata* Willd. | Fabaceae | Native | 2 |
| *Inga vera* Willd. | Fabaceae | Native | 6 |
| *Jacaranda caroba* (Vell.) DC. | Bignoniaceae | Endemic | 1 |
| *Jacaratia spinosa* (Aubl.) A.DC. | Caricaceae | Native | 1 |
| *Luehea divaricata* Mart. | Malvaceae | Native | 2 |
| *Mabea fistulifera* subsp. *fistulifera* Mart. | Euphorbiaceae | Endemic | 2 |
| *Maclura tinctoria* (L.) D.Don ex Steud. | Moraceae | Native | 2 |
| *Mimosa bimucronata* (DC.) Kuntze | Fabaceae | Native | 14 |
| *Mimosa* sp. L. | Fabaceae | Native | 1 |
| *Peltophorum dubium* (Spreng.) Taub. | Fabaceae | Native | 34 |
| *Psidium guajava* L. | Myrtaceae | Naturalized | 5 |
| *Pterogyne nitens* Tul. | Fabaceae | Native | 6 |
| *Sapindus saponaria* L. | Sapindaceae | Native | 2 |
| *Sapium haematospermum* Müll.Arg. | Euphorbiaceae | Native | 2 |
| *Schinus terebinthifolia* Raddi | Anacardiaceae | Native | 7 |
| *Schizolobium parahyba* (Vell.) Blake | Fabaceae | Native | 4 |
| *Spodias tuberosa* Arruda | Anacardiaceae | Nativa | 1 |
| *Spondias mombin* L. | Anacardiaceae | Native | 2 |
| *Syagrus romanzoffiana* (Cham.) Glassman | Arecaceae | Native | 1 |
| *Syzygium cumini* (L.) Skeels | Myrtaceae | Naturalized | 1 |
| *Tabebuia roseoalba* (Ridl.) Sandwith | Bignoniaceae | Native | 2 |
| *Tabernaemontana hystrix* Steud. | Apocynaceae | Native | 1 |
| *Talipariti pernambucense* (Arruda) Bovini | Malvaceae | Native | 3 |
| *Tecoma stans* (L.) Juss. Ex Kunth | Bignoniaceae | Naturalized | 9 |
| *Triplaris americana* L. | Polygonaceae | Native | 2 |
| *Zeyheria tuberculosa* (Vell.) Bureau ex Verl. | Bignoniaceae | Native | 11 |

**Table S2.** The perception of ecosystem services and disservices of farmers who managed and abandoned management of the main crop. Shown are the proportion farmers that answered “yes” for a given service or disservice, separately for farmers who managed and abandoned management of the main crop, as well as the P-value of a Fisher’s exact test. P-values lower than 0.05 indicate a significant difference between the two groups, whereas P-values lower than 0.1 indicate a trend.

|  | | **Proportion of ‘Yes’ answers from farmers who still manage**  **(n = 10)** | **Proportion of ‘Yes’ answers from farmers who abandoned management**  **(n = 20)** | **P** |
| --- | --- | --- | --- | --- |
| **Services** | Fruit collection | 0.8 | 0.6 | 0.42 |
|  | Honey collection | 0.4 | 0.25 | 0.43 |
|  | Medicinal herb collection | 0.4 | 0.2 | 0.38 |
|  | Firewood collection | 0.6 | 0.25 | 0.11 |
|  | Timber collection | 0.6 | 0.15 | 0.03^*^ |
|  | Aesthetical use | 0.8 | 0.4 | 0.06^†^ |
|  | Leisure use | 0.3 | 0.35 | 1 |
|  | Climatization use | 0.5 | 0.55 | 1 |
|  | Protection against the elements use | 0.4 | 0.45 | 1 |
|  | Shadow use | 0.9 | 0.85 | 1 |
|  | Pollinator presence | 0.9 | 1 | 0.33 |
|  | Bird presence | 1 | 1 | 1 |
|  | Wild sapling presence | 0.5 | 0.35 | 0.46 |
|  | Soil quality | 1 | 0.75 | 0.14 |
| **Disservices** | Venomous arachnid presence | 0.6 | 0.25 | 0.11 |
|  | Venomous snake presence | 0.5 | 0.55 | 1 |
|  | Small vertebrate presence | 0.9 | 0.85 | 1 |
|  | Large vertebrate presence | 0.5 | 0.15 | 0.08^†^ |
|  | Pest presence | 0.9 | 0.85 | 1 |

^*^ Significant
^†^ Trend

**Table S3.** Statistical overview of multiple regression models of biodiversity indices and structural characteristics as a function of abandonment of management, time since establishment, time since abandonment and the interaction term between time since establishment and time since abandonment. Shown are estimates of coefficients, chi-squared, degrees of freedom and P-values. We used backward model selection with a threshold of P < 0.1 to retain predictor variables to reach a final model and assessed significance using the function *Anova* in the package *car* (Fox et al., 2024).

|  | **Explanatory variables** | | | | | | | | | | | | | | | |
| --- | --- | --- | --- | --- | --- | --- | --- | --- | --- | --- | --- | --- | --- | --- | --- | --- |
|  | **Abandonment of management** | | | | **Time since establishment** | | | | **Time since abandonment** | | | | **Time since establishment × Time since abandonment** | | | |
| **Response variable** | Coeff. | Χ² | *D.f.* | P | Coeff. | Χ² | *D.f.* | P | Coeff. | Χ² | *D.f.* | P | Coeff. | X² | *D.f.* | P |
| Species richness | - | - | - | - | - | - | - | - | -0.03 | 2.24 | 1 | 0.13 | - | - | - | - |
| Shannon’s diversity index | - | - | - | - | - | - | - | - | -0.03 | 0.61 | 1 | 0.08^†^ | - | - | - | - |
| Abundance (DBH > 1cm) | -0.19 | 0.53 | 1 | 0.46 | - | - | - | - | - | - | - | - | - | - | - | - |
| Abundance (DBH < 1cm) | - | - | - | - | - | - | - | - | 0.03 | 0.19 | 1 | 0.66 | - | - | - | - |
| Evenness | - | - | - | - | - | - | - | - | -0.02 | 0.55 | 1 | 0.08^†^ | - | - | - | - |
| Canopy cover | - | - | - | - | 0.008 | 1.28 | 1 | 0.26 | - | - | - | - | - | - | - | - |

* Significant
^†^ Trend
- Removed through backward model selection

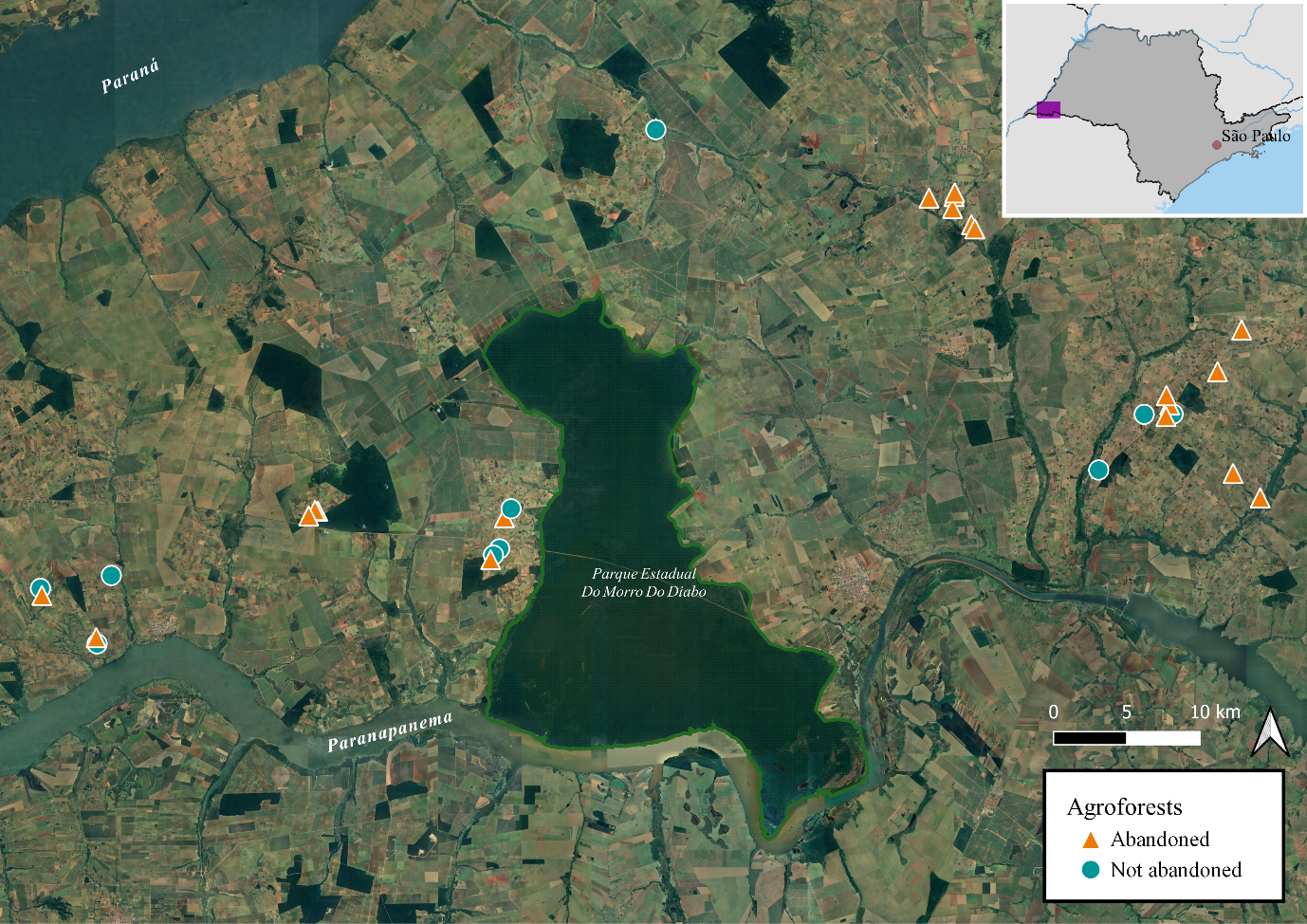

**Fig. S1.** Map of the region of *Pontal do Paranapanema* with the location of the thirty farms. Agroforests which were still maintained (n = 21) are illustrated as blue circles, and those that were abandoned (n = 20) as orange triangles. The largest remnant of Brazilian Atlantic Forest in this region is the *Morro do Diabo* State Park, which can be recognized as the dark green area with a green outline in the middle of the map. The inset in the top right shows the location of the region of *Pontal do Paranapanema* (purple square) within the state of São Paulo in southeastern Brazil.

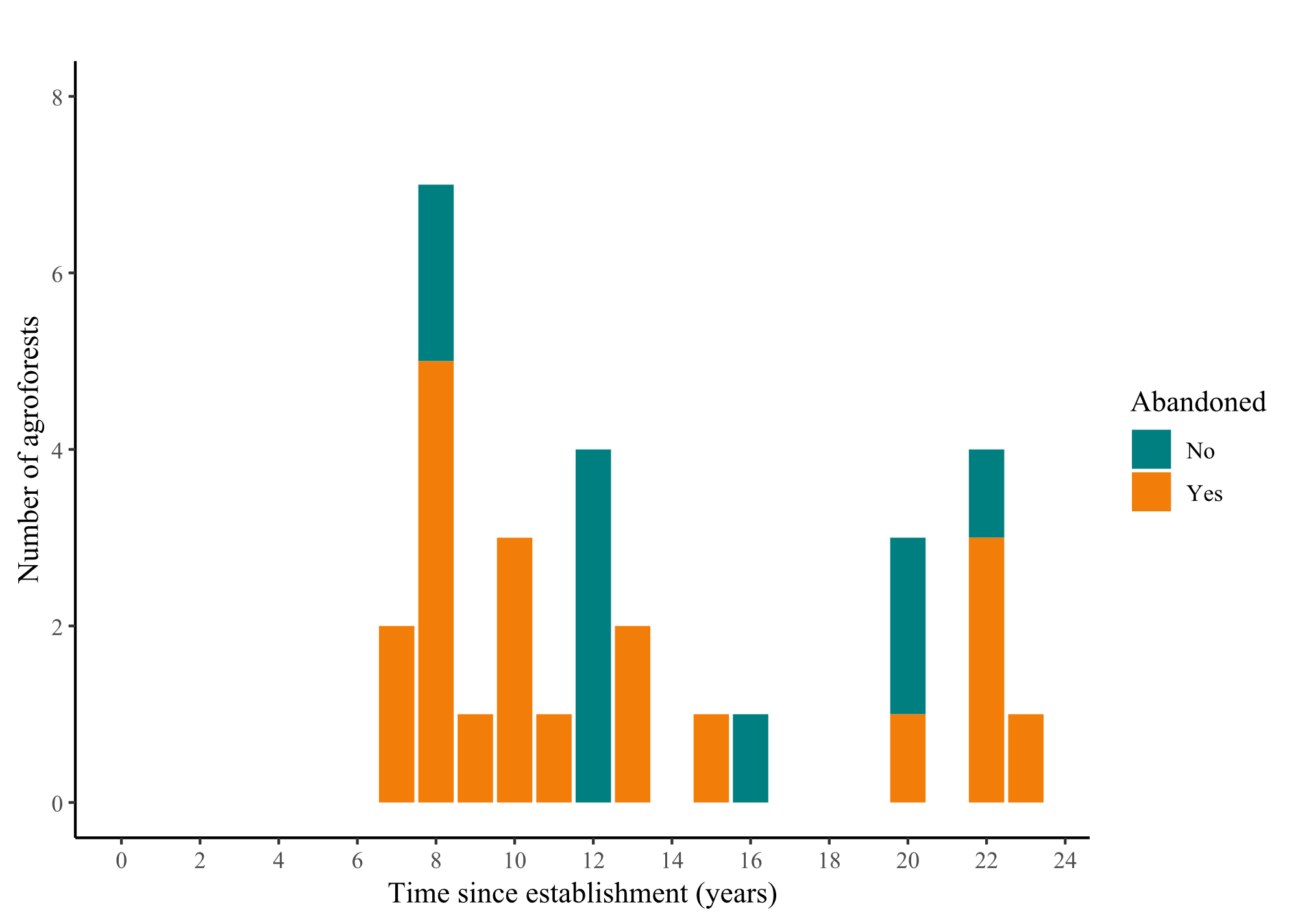

**Fig. S2.** Histogram showing the frequency distribution of time since establishment of thirty agroforests in southeastern Brazil. Turquoise represents agroforests which are still managed, and orange represent agroforests in which the management was abandoned. There is no difference in the time since establishment between managed and abandoned agroforests (t-test: t_20_ = 0.73, P = 0.47).

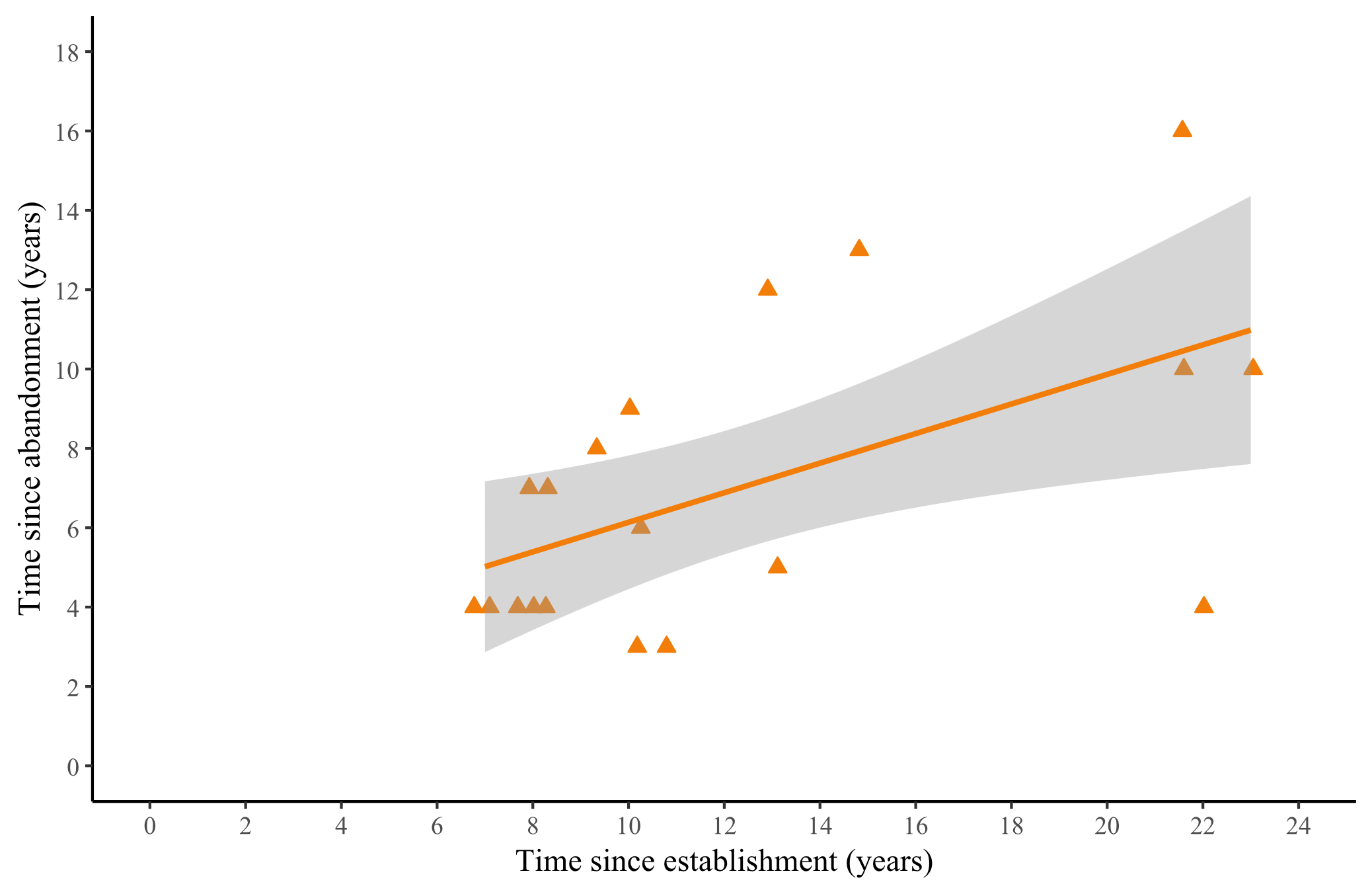

**Fig. S3.** The relationship between time since establishment and time since abandonment for agroforests where management of the main crop was abandoned. The dashed orange trendline is significant (r_27_ = 5.45, P = 0.01). Points are slightly jittered along the x-axis to avoid overlap.

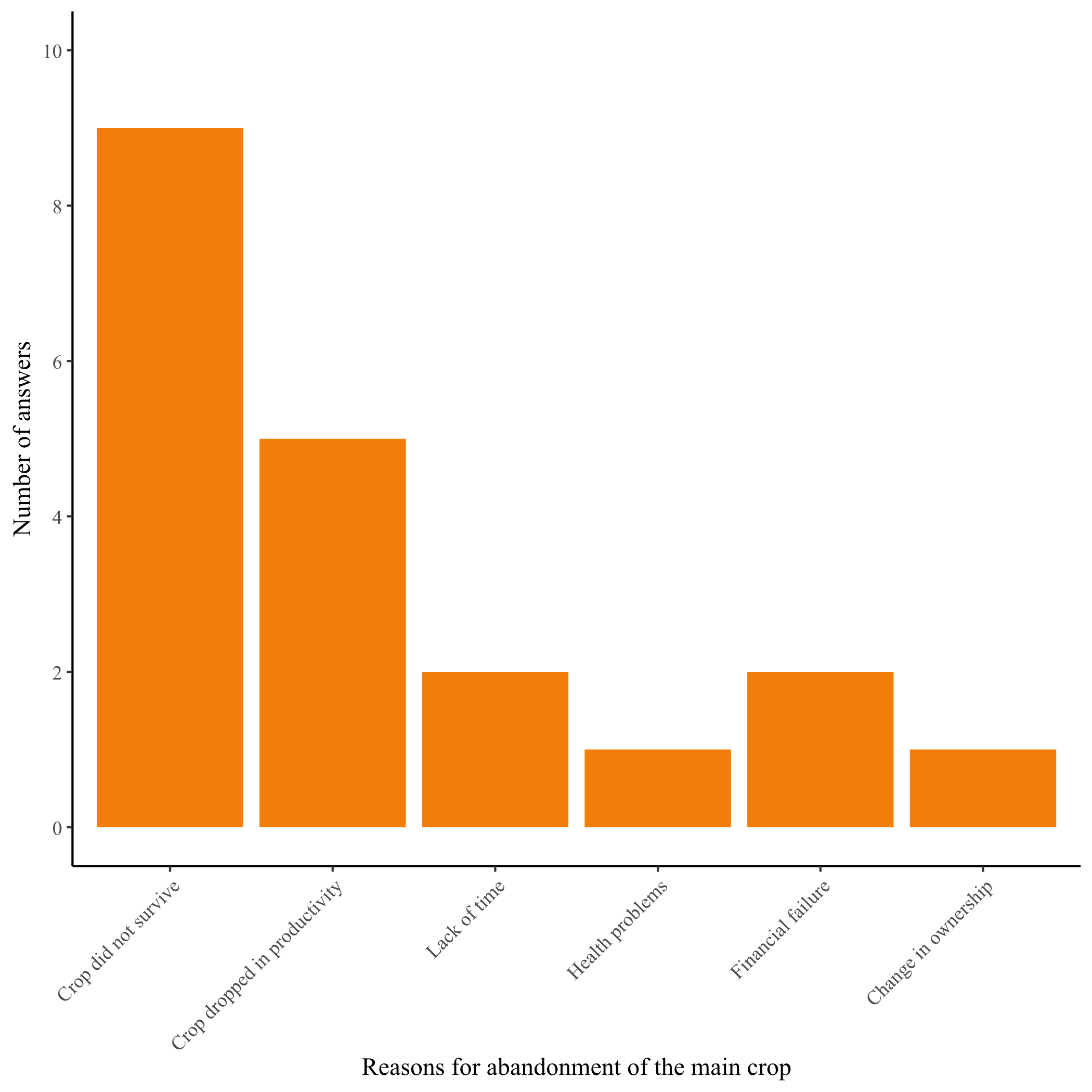

**Fig. S4.** Frequency distribution of answers farmers (n = 20) gave when asked why they abandoned management of the main crop. In case of financial failure, the farmer did not consider the continuation of management financially viable.

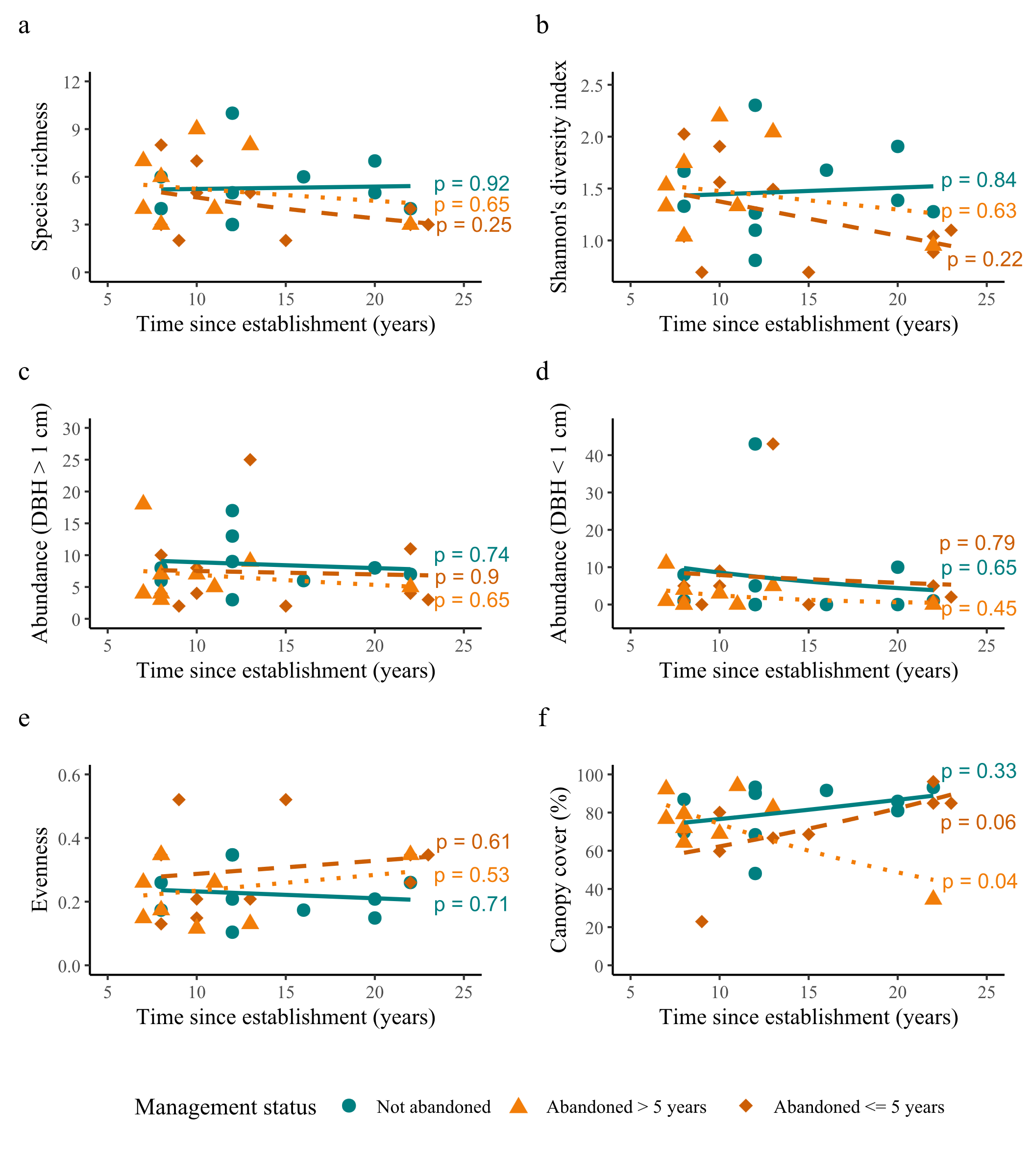

**Fig. S5.** Biodiversity and structural indexes as a function of time since establishment and time since abandonment. Turquoise circles represent agroforests which are still managed, orange triangles represent agroforests where management of main crop was abandoned five or fewer years ago and red diamonds represent agroforest which management of main crop was abandoned more than 5 years ago. Presented are (a) species richness, (b) Shannon’s diversity index, (c) abundance of woody plants with DBH larger than 1 cm, (d) abundance of woody plants with DBH smaller than 1 cm, (e) evenness and (f) canopy cover.
