## Appendix A for "Open-land-derived agroforestry and effects of abandonment of management of the main crop on ecosystem services and woody plant diversity"

Appendix A – Interview

1. Type: ☐ Managed	☐ Abandoned
2. What was your land used for before you settled in?
3. When did you settle in?
4. When did you establish your agroforest?
5. Did you have any help when implementing agroforestry? If yes, from whom and what kind of help?

☐ Financial	☐ Resources	☐ Work force

☐ Other:________________________________________________________

6. What was the area of agroforestry used for before establishing it?

☐ Pasture	☐ Cropland	☐ Wild secondary vegetation

☐ Other:_________________________________________________________

7. What was the reason for the choice of location for establishing the agroforest?
8. Do you regret the adoption of the agroforestry system?
9. What was the initial main product cultivated for selling in the agroforest? What about for consumption?
10. Did you do any kind of management during the establishment of the agroforest?

☐ Watering	☐ Fertilizing

☐ Mulching	☐ Mowing

☐ Hand weeding	☐ Soil overturning

☐ Thinning of shrubs/saplings	☐ Cattle integration

☐ Other:_________________________________________________________

11. How many hours per week do/did you use to spend managing it?
12. Did you change the intensity of management over time?
13. Do you still do any kind of management currently? If yes, which?

☐ Watering	☐ Fertilizing

☐ Mulching	☐ Mowing

☐ Hand weeding	☐ Soil overturning

☐ Thinning of shrubs/saplings	☐ Cattle integration

☐ Other:_________________________________________________________

14. (For abandoned agroforests) When did you cease to manage it?
15. (For abandoned agroforests) What was the main reason that led you to cease the management?

☐ Financial failure	☐ Lack of institutional support

☐ Costs	☐ Health issues

☐ Loss in productivity	☐ Crop failure

☐ Other:__________________________________________________________

16. (For abandoned agroforests) If you still do management, why do you do it?
17. What are notable benefits you get from the agroforest?

☐ Fruits. Which?____________________________________________________
(In the past few years there was: ☐ Increase, ☐ No Change, ☐ Decrease)

☐ Medicinal Plants. Which?___________________________________________
(In the past few years there was: ☐ Increase, ☐ No Change, ☐ Decrease)

☐ High soil quality
(In the past few years there was: ☐ Improvement, ☐ No Change, ☐ Deterioration)

☐ Beehives
(In the past few years there was: ☐ Increase ☐ No Change ☐ Decrease)

☐ Firewood	☐ Timber	☐ Relaxation, cultural, or religious use

☐ Pleasant aesthetics	☐ Climatization/Shade

☐ Other:___________________________________________________________

18. Are there any kind of problems that you feel come from having an Agroforest in your property?

☐ Venomous arachnids (spiders/scorpions). Which?________________________________________________________
(In the past few years there was: ☐ Increase ☐ No Change ☐ Decrease)

☐ Snakes. Which?_______________________________________________
(In the past few years there was: ☐ Increase ☐ No Change ☐ Decrease)

☐ Pests. Which?________________________________________________
(In the past few years there was: ☐ Increase ☐ No Change ☐ Decrease)

☐ Large predator. Which?_________________________________________
(In the past few years there was: ☐ Increase ☐ No Change ☐ Decrease)

☐ Other:_______________________________________________________

19. I am interested in other aspects of biodiversity, could you tell me if you notice:

☐ Wild saplings you did not plant yourself.
(In the past few years there was: ☐ Increase ☐ No Change ☐ Decrease)

☐ Pollinators, such as bees and butterflies.
(In the past few years there was: ☐ Increase ☐ No Change ☐ Decrease)

☐ Birds. Which?________________________________________________
(In the past few years there was: ☐ Increase ☐ No Change ☐ Decrease)

☐ Small rodents. Which?__________________________________________
(In the past few years there was: ☐ Increase ☐ No Change ☐ Decrease)
